# Expanding the CarD interaction network: CrsL is a novel transcription factor in *Mycobacterium smegmatis*

**DOI:** 10.1101/2024.12.02.626298

**Authors:** Mahmoud Shoman, Jitka Jirát Matějčková, Marek Schwarz, Martin Černý, Nabajyoti Borah, Viola Vaňková Hausnerová, Michaela Šiková, Hana Šanderová, Petr Halada, Martin Hubálek, Věra Dvořáková, Martin Převorovský, Jana Holubová, Ondřej Staněk, Libor Krásný, Lukáš Žídek, Jarmila Hnilicová

## Abstract

Bacterial transcription regulation is critical for adaptation and survival. CarD is an essential transcription factor in mycobacteria involved in regulation of gene expression. We searched for CarD interaction partners in the model organism *Mycobacterium smegmatis* and identified two proteins: ApeB (MSMEG_5828) and an uncharacterized protein, which we named CrsL (MSMEG_5890). While ApeB interacted with CarD only when CarD was overexpressed, CrsL associated with CarD at its physiological levels. CrsL is a 5.7 kDa protein shown by NMR to be intrinsically disordered. CrsL homologs are present in actinobacteria including pathogenic species such as *Mycobacterium tuberculosis*. CrsL directly interacts with CarD and binds RNAP. ChIP-seq showed that CrsL associates with promoters of actively transcribed genes and ∼75 % of these regions are also associated with CarD. RNA-seq showed ∼50% and ∼66% overlap in differentially expressed genes between CrsL and CarD knockdowns during exponential and stationary phases, respectively. CrsL represses expression of DesA desaturase (*MSMEG_5773*) and DEAD/DEAH-box RNA helicase *MSMEG_1930*, which are important for adaptation to cold stress. Furthermore, CrsL promotes the growth of *M. smegmatis* at elevated temperature. In summary, this study identifies CrsL as a novel actinobacterial transcription factor and provides a basis for its further investigation.

## INTRODUCTION

Mycobacteria include not only slow-growing species like *Mycobacterium tuberculosis*, responsible for tuberculosis, but also many rapidly growing nontuberculous mycobacteria that can cause a variety of infections. Certain nontuberculous mycobacteria are capable of growth in a wide range of temperatures, such as in the water circuits of heater-cooler units, medical devices that control the patient’s body temperature during open-heart surgery. *Mycobacterium chelonae* was reported to infect patients during cardiac surgeries who were exposed to contaminated aerosols from these units (1,2). To survive varying temperatures and environmental challenges, mycobacteria, like other bacteria, rely on transcriptional regulation. Here, we studied transcriptional regulation in *Mycobacterium smegmatis*, a well-established model to study these regulatory mechanisms in mycobacteria.

Bacterial transcription involves the synthesis of RNA from a DNA template, a process mediated by a single type of DNA-dependent RNA polymerase (RNAP) (3). The RNAP core consists of several subunits (α2ββ′ω) (4). These subunits associate with a sigma (σ) factor to form the RNAP holoenzyme, that can recognize promoter sequences and initiate transcription (5–7). All bacteria have one primary σ factor (7,8). The primary σ factor in mycobacteria is σ^A^ and is essential for bacterial growth (9,10). Besides σ^A^, the mycobacterial alternative σ factor σ^B^ recognizes similar promoter sequences as σ^A^, and consequently, regulons of these σ factors partly overlap (11–14). In addition, σ^B^ regulates stress-responsive genes (15–17).

The mycobacterial transcription machinery requires additional transcription factors − RbpA and CarD, which are not present in *Escherichia coli* (18,19). These factors are essential global regulators both in *M. smegmatis* and *M. tuberculosis* (20–24).

RbpA consists of four domains and binds to RNAP containing σ^A^ or σ^B^ but not the other alternative σ factors (20,25–27). It assists in promoter unwinding and formation of the catalytically active open complex (11,23). RbpA was proposed to modify the structure of the RNAP core, increasing the competitiveness of the σ^A^ over the alternative σ factors (27). Additionally, RbpA has been suggested to play a role in the release of rifampicin from RNAP and is also involved in the stress response (22,27–30).

CarD consists of two domains. The CarD N-terminal domain has a conserved structure and interacts with RNAP (RNAP interaction domain, RID), while the C-terminal domain binds to DNA (DNA-binding domain, DBD) (31). CarD affects formation/stability of the open complex in a promoter dependent manner (32). CarD acts as a global transcription regulator (31–35) and regulates many genes including ribosomal RNAs encoding genes (33,36). The mycobacterial growth is altered when CarD function is disrupted by mutations targeting the RNAP interaction domain (RID), the DNA-binding domain (DBD), or a conserved tryptophan residue (W85) (36). Furthermore, weakening the mycobacterial RNAP-CarD interaction results in cells being more sensitive to stress conditions, including oxidative stress, DNA damage, and the effect of some antibiotics (37,38). The level of CarD decreases during stationary phase and starvation. This decrease is mediated by *carD* antisense RNA and Clp protease (38).

Apart from CarD and RbpA, mycobacteria contain a unique ∼300 nt long Ms1 RNA (39). Ms1 RNA is highly expressed during stationary phase of *M. smegmatis* growth and interacts directly with the RNAP core without any σ factor and regulates the RNAP level in *M. smegmatis* (40,41). Ms1 has homologs in various actinobacterial species (42), including MTS2823 RNA, that is highly expressed in *M. tuberculosis* (43) and interacts with RNAP (44).

In this study, we sought to expand our knowledge of the mycobacterial transcription machinery and searched for CarD interaction partners in *M. smegmatis.* We identified two novel interaction partners of CarD – ApeB and CrsL. While ApeB interacts with CarD when CarD level is elevated, CrsL binds to CarD also when the CarD level is unchanged. Therefore, the main subsequent focus was on CrsL. We first validated the CrsL-CarD interaction and showed that both can associate with RNAP. We then characterized the structure of CrsL, defined its binding sites over the chromosome and determined effects of CrsL depletion on the bacterial growth and the transcriptome. Taken together, our data suggests that CrsL is a novel mycobacterial transcription factor.

## MATERIALS AND METHODS

### Construction of the bacterial strains

The detailed descriptions of all bacterial strains and oligonucleotides sequences used for strain construction are listed in the Supplementary Data.

#### CrsL-6xHis strain

The CrsL-6xHis (at its N-terminus) was generated by PCR cloning using Q5 High-Fidelity DNA Polymerase (NEB) with primers #4539 + #4540 and *M. smegmatis* genomic DNA (*LK2980*) as template. The PCR product was cloned into pET22b plasmid via *NdeI*/*XhoI* restriction sites resulting in *LK3496* (*E. coli* DH5α), verified by sequencing and transformed into *E. coli* DE3 cells, resulting in LK3499 strain. The CarD-NT expression strain (LK3209) was constructed previously (45).

#### ApeB-FLAG and CrsL-FLAG strains

The ApeB (*MSMEG_5828*) and CrsL (*MSMEG_5890*) genes were amplified by PCR using Q5 High-Fidelity DNA Polymerase (NEB) with primers #3415 + #3417 (for ApeB), and #3909 + #3910 (for CrsL) and *M. smegmatis* genomic DNA as template. The C-terminal 1× FLAG-tag (DYKDDDDK) was encoded within the reverse PCR primers. CrsL-FLAG was cloned into pTetInt integrative plasmid (46) via *NdeI*/*HindIII* restriction sites and verified by sequencing. The plasmids were integrated into *M. smegmatis* mc^2^ 155 genome by electroporation resulting in ApeB-FLAG (*LK2767*) and CrsL-FLAG (*LK3051*) strains. The CarD-FLAG (*LK2539*), RbpA-FLAG (*LK2541*), HelD-FLAG (*LK2589*) strains were generated previously (45,47).

#### ApeB-gFLAG and CarD-gFLAG strains

The strains with the FLAG-tagged in the genome (native locus) − ApeB-gFLAG (*LK2846*) and CarD-gFLAG (*LK2899*) were designed using NEBuilder Assembly Tool (https://nebuilder.neb.com/) and constructed using Gibson assembly kit (NEB) as following. The cassettes were generated by assembling three PCR fragments amplified with Q5 High-Fidelity DNA Polymerase (NEB); i) 500 bp long fragment [left arm, LA] homologous to the 3’ terminal part of ApeB (*MSMEG_5828*) or CarD (*MSMEG_6077*) containing the FLAG-tag (DYKDDDDK) amplified by primer #3378 + #3379 or #3372 + #3373, respectively; ii) 500 bp long fragment [right arm, RA] homologous to the 5’ end of the ApeB or CarD gene amplified by primer #3382 + #3383 or #3376 + #3377, respectively; iii) The left and right arms of ApeB and CarD were flanked by hygromycin resistance cassette amplified from *LK1463* strain by primer #3380 + #3381 or #3374 + #3375, respectively. The three fragments (LA, HYG, RA) of ApeB or CarD were assembled into pUC18 (*LK222*) plasmid with *HindIII*/*XbaI* or *HindIII*/*BamHI*, respectively, using Gibson assembly Cloning kit (NEB). The Gibson assembly constructs were transformed into *E. coli* DH5α. The resulting ApeB (*LK2796*) and CarD (*LK2795*) strains were verified by sequencing. The fragments encompassing the cassettes were subsequently transformed into the *M. smegmatis* pJV53 (*LK1321*) or *M. smegmatis* pYS1 (*LK2713*) strains for homologous recombination and individual clones were selected, resulting in chromosomal FLAG-tagged gene encoding ApeB-gFLAG (*LK2846*) and CarD-gFLAG (*LK2899*) strains.

#### carD and crsL CRISPR depletion strains

The sgRNA oligonucleotides are listed in Supplementary Data. The sgRNA targeting either the gene (*crsL*) or the gene promoter (*carD*) were cloned into PLJR962 according to the previously established protocol (48), verified by sequencing and transformed by electroporation into *M. smegmatis* wild type (*LK2598)* resulting in strains *LK3301* (*MSMEG_6077*, *carD* sgRNA) and *LK3302* (*MSMEG_5890*, *crsL* sgRNA). The negative control non-targeting control sgRNA strain (*nc*WT, *LK2261*) and *LK2203* (*MSMEG_2758*, *sigA* sgRNA) were generated previously (48–50).

### Bacterial growth conditions

*M. smegmatis* mc^2^ 155 strains were grown onto Middlebrook 7H10 (Difco) for 2-3 days at 37 °C. When required, the media was supplemented with antibiotics: kanamycin (20-25 μg/mL) or hygromycin (50 μg/mL). The strains were inoculated into overnight culture in Middlebrook 7H9 media (Difco) supplemented with 0.2% glycerol and 0.05% Tween 80 at 37 °C. The overnight cultures were then diluted into OD_600_ 0.1 and allowed to grow for exponential phase (OD_600_ ∼0.5; 6 h of growth) or early stationary phase (OD_600_ ∼2.5–3; 24 h of growth).

#### FLAG-tagged and CRISPR knockdown strains

The strains were inoculated to OD_600_ 0.1. For exponential phase samples (6 h growth, OD_600_ ∼0.5), anhydrotetracycline (ATc) was added 3 h after inoculation in different concentrations: 100 ng/mL (for CRISPR strains), 10 ng/mL (for ChIP-seq and IP), 25 ng/mL (for overexpression) or 1 ng/mL (for CarD-FLAG optimization) and cells harvested after additional 3 h. For stationary phase (24 h growth, OD_600_ ∼2.5-3), ATc was added 8 h after inoculation and cells harvested after 16 h later. For CarD-FLAG optimization, the ATc (1 ng/mL) was added after 21 h and cells harvested after 3 h later. The discs containing 4 μl of 100 ng/mL of ATc were placed on 7H10 plates with an equal volume (1 mL) of the strains (OD_600_ ∼1) and incubated at 37 °C for 2-3 days.

For the growth curves at elevated temperature, overnight cultures of ncWT and *carD* knockdown strains were initially grown at the optimal temperature (37°C) with ATc (100 ng/mL) for approximately 16 hours to deplete CrsL. They were then diluted to OD_600_ ∼0.1 in 7H9 media with ATc (100 ng/mL). 6 h after inoculation, the temperature was raised to 45°C and cells were cultivated for ∼35 h. The cells were grown in BioSan RTS-8 Multi-Channel Bioreactor and OD_600_ was measured throughout the growth.

### Immunoprecipitation (FLAG pulldown)

100 mL of *M. smegmatis* exponential and stationary phase cells were pelleted and washed in lysis buffer [20 mM Tris–HCl pH 7.9, 150 mM KCl, 1 mM MgCl_2_] and pelleted again and stored at –70°C. The pellets were resuspended in 3 mL of Lysis buffer supplemented with phenylmethylsulfonyl fluoride (PMSF) and Protease inhibitor cocktail [20 mM Tris–HCl pH 7.9, 150 mM KCl, 1 mM MgCl_2_, 1 mM dithiothreitol (DTT), 0.5 mM PMSF, Protease Inhibitor Cocktail Set III protease inhibitors (Calbiochem)], sonicated 15 × 10 s with 1 min pauses on ice and centrifuged at 8960 × g for 15 min at 4°C. Equal amount of cell lysates (2–4 mg of proteins) from the FLAG-tagged strains were incubated for 16–18 h overnight at 4°C with 25 μl of M2 anti-FLAG resin (Sigma Aldrich). The captured protein complexes with agarose gel beads were washed 4× with 0.5 mL of lysis buffer. FLAG-tagged proteins were eluted by 60 μl of 3× FLAG Peptide (Sigma F4799) diluted in Tris-buffer saline (TBS: 50 mM Tris-HCl pH 7.5, 150 mM NaCl) to a final concentration of 150 ng/mL. Proteins with the captured complexes were resolved on SDS-PAGE gels (Nu-PAGE, 4–12% Bis–Tris precast gels, Invitrogen) and stained with SimplyBlue SafeStain (Invitrogen), or stained with silver (Pierce Silver Stain kit for Mass Spectrometry or SilverQuest Silver Staining Kit (Thermo Fisher Scientific) and/or analyzed by western blotting. The identity of the protein bands was determined by MALDI-FTICR mass spectrometry as described previously (47).

### CrsL-His protein purification for *in vitro* assay and antibody production

*E. coli* strains used for protein purification, CarD-NT (*LK3209*) and CrsL-His (*LK3499*), were inoculated into overnight cultures in LB medium at 37 °C. The overnight cultures were then diluted to OD_600_ = 0.03 into the LB medium with ampicillin (100 μg/mL) and shaken at 120 rpm at 37 °C for ∼ 3 h (until OD_600_ ∼0.6); then the expression was induced with 0.8 mM IPTG. The temperature was then reduced to room temperature (∼20°C) and shaken for an additional 3 h. The cultures were cooled and centrifuged using Beckman Coulter JA-10 rotor at 4°C, 6000 x g for 10 min, The pellets were resuspended in 1x T-buffer (300 mM NaCl, 50 Mm Tris-HCl pH 7.5, 5% glycerol) and centrifuged 5400 x g for 10 min and the pellets were stored in -80°C. The pellets were sonicated (Sonopuls HD3100, Bandelin [Germany]; 50 % amplitude, 15 x 10 s pulse, 1 min pause on ice) in 1x T-buffer (supplemented with 3 mM β-mercaptoethanol). The samples were centrifuged at 27000 x g, 4°C (Beckman Coulter, JA 25-50 rotor) for 10 min and the cell lysates were incubated with 1 mL of prewashed Ni-NTA Agarose beads (QIAGEN) shaking at 4°C for 1.5 h. The samples were centrifuged at 2000 rpm, 4°C for 5 min and Ni-NTA beads were resuspended in 10 mL 1x-T buffer with and centrifuged again at 2000 rpm, 4°C for 5 min. The Ni-NTA beads were resuspended in 10 mL 1x-T Buffer (containing 30 mM imidazole) and centrifuged again at 2000 rpm, 4°C for 5 min. The Ni-NTA beads were then resuspended 500 μl of 1x-T Buffer (400 mM imidazole) and serial fractions of the eluted proteins were collected, and their concentrations and purity were measured by Bradford assay and on SDS-PAGE, respectively. The selected fractions containing CrsL-His protein that had the highest concentration and the highest purity were pooled and dialyzed for 20 h against Buffer A (50 mM Tris-HCl, pH 8, 100 mM NaCl, 0.5 mM EDTA, 0.5 mM DTT, 5% glycerol) in Slide-A-Lyzer Dialysis Cassette 2,000 MWCO (Thermo Fisher Scientific). Protein samples and the eluent were visualized and checked by SDS-PAGE gels stained with SimplyBlue SafeStain (Invitrogen). Protein fractions were mixed and dialyzed against storage buffer (50 mM Tris-HCl, pH 8, 100 mM NaCl, 50% glycerol, 3 mM β-mercaptoethanol) and stored at -20°C.

For CarD-NT protein, the overexpression protocol was similar to above with additional TEV protease cleavage as described previously (45). Both CrsL-His and CarD-NT proteins were subsequently subjected to additional purification via ӒKTA pure™ chromatography system (Superdex 75 column) and eluted with Buffer B (50 mM Tris-HCl pH 8, 100 mM NaCl, 5% glycerol, 3 mM β-mercaptoethanol). CrsL-His protein was then used for mice immunization and anti-CrsL antibody production as described in the following section.

### Animal experiments and immunizations

All animal experiments were approved by the Animal Welfare Committee of the Institute of Molecular Genetics of the Czech Academy of Sciences, v. v. i., in Prague, Czech Republic. Handling of animals was performed according to the Guidelines for the Care and Use of Laboratory Animals, the Act of the Czech National Assembly, Collection of Laws no. 246/1992. Permission no. 19/2020 was issued by the Animal Welfare Committee of the Institute of Molecular Genetics of the Czech Academy of Sciences in Prague.

Five-week-old female BALB/cByJ mice (Charles River, France) were immunized by intraperitoneal injection with CrsL protein (20 μg in 200 μl) adjuvanted with aluminum hydroxide (Alum, SevaPharma, Czech Republic). Mice received three doses of the protein in 2 weeks interval and 1 week after the third immunization, the blood was collected from anesthetized animals (i.p. injection of 80 mg/kg ketamine and 8 mg/kg xylazine) by retroorbital puncture method. Sera were recovered from the supernatant after centrifugation of clogged blood at 5000 x g for 10 min at 8 °C and stored at –20 °C. The specificity of the antibody was checked with 1 μg of *M. smegmatis* cell lysate with 1:5000 antibody dilution.

### Western blotting and FAR-western blotting

The samples were resolved on SDS-PAGE gels and detected by western blotting using anti-RNAP β subunit antibody [clone 8RB13] (BioLegend) or anti-σ^70^ antibody [clone 2G10] (BioLegend), anti-CarD antibody, anti-RbpA antibody, anti-CrsL antibody, anti-GroEL [5177] antibody (Santa Cruz Biotechnology) and a HRP-labeled anti-mouse IgG antibody (Sigma-Aldrich). The blots were incubated with SuperSignal West Pico PLUS Chemiluminiscent substrate (Thermo Fisher Scientific) and the signals were detected on film with different exposition time.

FAR-western blotting was performed according to the previously established protocol (51). Briefly, BSA, CarD-NT, CrsL-His proteins were loaded in duplicate and resolved on SDS-PAGE gels and blotted onto Amersham Protran Nitrocellulose Membrane (Sigma-Aldrich). The membranes were incubated with decreasing concentrations of guanidine-HCl (Thermo Fisher Scientific) for protein denaturation and renaturation. The membranes were blocked with 5% milk and washed with PBST buffer. The membranes were cut into two parts. The first part (negative control) was incubated directly with anti-CarD antibody. The second part was incubated first with purified CarD protein (20 μg in 10 mL) overnight to allow CrsL-CarD complex formation and then with anti-CarD antibody. The membranes were washed with PBST and incubated with HRP-labeled anti-mouse secondary antibody (Sigma-Aldrich) and the signals were detected.

### ChIP-seq

CrsL-FLAG ChIP-seq experiments were performed in parallel with HelD, RbpA and CarD ChIP-seq (45). Briefly, 2 mg of protein cell lysates were incubated with 20 μl of M2 anti-FLAG resin (Sigma Aldrich). The captured complexes were washed twice with RIPA buffer (150 mM NaCl, 1% Triton X-100, 0.5% deoxycholate, 0.1% SDS, 50 mM Tris-HCl pH 8.0, 0.5 mM EDTA), four times with LiCl buffer (100 mM Tris-HCl, pH 8.5, 500 mM LiCl, 1% Triton X-100, 1% deoxycholate), two times with RIPA and twice with TE buffer (10 mM Tris-HCl pH 8.0, 1 mM EDTA). Protein-DNA complexes were eluted with elution buffer (50 mM Tris-HCl pH 8, 0.66 mM EDTA, 1 % SDS) for 10 min at 65 °C, decrosslinked in the presence of 200 mM NaCl for 5 h at 65 °C, treated with 100 µg/mL RNase A for 1 h at 37 °C and 400 µg/mL proteinase K for 30 min at 45 °C. DNA was purified with the QIAGEN PCR purification kit and eluted with 100 μl of Elution Buffer. 40 µl of immunoprecipitated DNA sample or 10 ng of DNA input were used for library construction according to the NEXTFLEX ChIP-Seq Kit manual including the Size-Selection Cleanup step B2. Pooled barcoded libraries (biological triplicates) were sequenced in single lanes using the Illumina NextSeq 500/550 High Output Kit v2 in 75 bp single end regime.

### RNA-seq

Before total RNA extraction, mRNA spike-in mix composed of 4 different eukaryotic mRNAs was added (Plat, Moc, Elav2 and nLuc; the mRNAs were prepared by *in vitro* transcription from pJET plasmids using the MEGAscript T7 Transcription Kit (Thermo Fisher Scientific), the sequence of the mRNAs is provided in the Supplementary Material). The amount of RNA spike-in was 10 ng per 30 mL of culture at an OD_600_ of 0.5. Each frozen cell pellet was resuspended in 240 μl TE buffer (pH 8.0) plus 60 μl LETS buffer (50 mM Tris-HCl pH 8.0, 500 mM LiCl, 50 mM EDTA pH 8.0, 5% SDS) and 600 μl acidic (pH∼3) phenol/chloroform (1:1). Samples were sonicated in a fume hood, centrifuged, the aqueous phase extracted two more times with acidic phenol/chloroform and precipitated with ethanol. RNA was dissolved in double distilled water and treated with DNase (TURBO DNA-free Kit, Invitrogen). 1 μg of DNase treated RNA was ribodepleted with riboPOOL Kit Pan-Actinobacteria (siTOOLs). The sample integrity was checked using Agilent 2100 Bioanalyzer Pico Chip. The ribodepleted RNA sample (20-100 ng) was used for library construction according to the NEXTFLEX Rapid Directional RNA-Seq Kit. The libraries were checked using Agilent 2100 Bioanalyzer Nano Chip. Pooled barcoded libraries were sequenced in single lane with Illumina NextSeq 500/550 High Output Kit v2 in 75 bp single end regime at Institute of Molecular Genetics AS CR, Prague, Czech Republic.

### RT-qPCR and northern blotting

5 μl RNA (∼ 2.5 μg) was reverse transcribed into cDNA (20 μl reaction, SuperScriptIII, Invitrogen) using random hexamers and amplified by RT-qPCR in a LightCycler 480 System (Roche Applied Science) in duplicate reactions containing LightCycler 480 SYBR Green I Master and 0.5 μM primers (each). Gene specific primers were designed with Primer3, sequences are listed in Supplementary Data.

Negative controls (no-template reactions and reactions with RNA as the template to check for genomic DNA contamination) were included in each experiment, the quality of the PCR products was determined by dissociation curve analysis and the efficiency of the primers determined by standard curves. The relative mRNA levels were quantified based on threshold cycles (Ct) for each PCR and normalized to the Plat mRNA spike-in value using the formula 2^(Ct^(spike)^ − Ct^(mRNA)^). The relative expression (E) was then normalized to the control strain (E =E_depleted_ /E_control_).

RNAs were resolved on a 7 M urea 7% polyacrylamide gel and transferred onto Zeta-Probe nylon membrane (Biorad) according to the protocol described in Panek et al. (39). 5′ biotinylated oligonucleotide probes (**Supplementary data**) were hybridized to the membrane and detected with Tropix CDP STAR substrate (ThermoFischer Scientific, Applied Biosystems) according to the manufacturer’s instructions.

### Glycerol gradient ultracentrifugation

*M. smegmatis* exponential and stationary phase cells were pelleted and resuspended in 20 mM Tris–HCl pH 8, 150 mM KCl, 1 mM MgCl_2_, 1 mM dithiothreitol (DTT), 0.5 mM phenylmethylsulfonyl fluoride (PMSF) and Calbiochem Protease Inhibitor Cocktail Set III protease inhibitors, sonicated 15 × 10 s with 1 min pauses on ice and centrifuged. Protein extracts (1 mg) were loaded on a linear 10–30% glycerol gradient prepared in gradient buffer (20 mM Tris–HCl pH 8, 150 mM KCl, 1 mM MgCl_2_) and fractionated by centrifugation at 130 000 × g for 17 h using an SW-41 rotor (Beckman). The gradient was divided into 20 fractions and proteins from each fraction were resolved on SDS-PAGE gels and detected by western blotting.

### [^13^C,^15^N] -double labeled CrsL protein purification for NMR

Expression cultures were grown in 2 L of minimal media (M9) containing 1M MgSO_4_, 500 mM CaCl_2_, 100 mM MnCl_2,_ 50mM ZnSO_4_ and 50 mM FeFl_3_ (52) and supplemented with ^15^NH_4_Cl and [^13^C] glucose. The cultures were incubated at 37 °C and shaken at 120 rpm for ∼ 3 h (until OD_600_ reached ∼0.6); expression was induced with 0.4 mM IPTG. The temperature was then reduced to room temperature (∼20°C) and the cultures were shaken for an additional 3 h. The protein was then purified according to the protocol mentioned in the previous section (protein purification). The [^13^C, ^15^N] uniformly labeled CrsL was dialyzed against the NMR buffer (20 mM sodium phosphate buffer, pH 7, 10 mM NaCl. 0.5 mM TCEP and 1 mM NaN_3_). The [^13^C,^15^N]-CrsL was additionally concentrated using Vivaspin® 15R Centrifugal Concentrator 2,000 MWCO (Sartorius).

### NMR measurements

All experiments were perfomed using a 950 MHz NMR spectrometer Bruker Avance III HD equipped with the TCI cryogenic probe head with z-axis gradients. For the measurements, the temperature was set to 27 °C and was calibrated according to the chemical shift differences of pure methanol peaks. Samples of 0.8 mM [^13^C,^15^N] labeled CrsL in NMR buffer were mixed with 10% Deuterium Oxide (D2O). For the backbone assignment, 0.15 mM uniformly labelled [^13^C,^15^N]-CrsL and the standard set of 3D triple-resonance experiments consisting of HNCA, HN(CO)CA, HNCACB, CBCA(CO)NH, and HNCO (53) was used. Spectra were processed using NMRPipe software (54) and the subsequent assignment of the peaks was done using the software Sparky 3.115 (T.D. Goddard and D. G. Kneller, University of California). Values of secondary structure propensity (SSP) were calculated with the SSP (1.0) script using chemical shifts of assigned backbone residues (55). For reference, we used predicted chemical shifts of the random coil form of our protein calculated from the sequence using Poulsen IDP/IUP random coil chemical shifts script (56–58). Neighbor corrected structural propensity (ncSP) values were computed using ncSP calculator (59), using chemical shifts based on Tamiola, Acar and Mulder (2010). Values corresponding to the AlphaFold2 predicted structure were calculated using the SHIFTX2 web tool (60).

### *in silico* predictions

Multiple sequence alignments were constructed based on hits from PHI-BLAST queries and results from DeepMSA (2.0) (61,62). We curated putative homologs based on similarity and compared them with KEGG Sequence Similarity DataBase (63). The final multiple sequence alignments was prepared using ClustalW using UGENE toolkit and plotted in ESPript (3.0) (64,65). The sequence logo was created using the WebLogo (3.7.12) software (66). The disorder was predicted using PSIPRED 4.0, NetSurfP 3.0, IUPred2A, ESpritz 1.3 and flDPnn predictors (67–71). In the case of IUPred2A, the IUPred2 short disorder was chosen. For predictions using ESpritz, we chose the NMR prediction type and Best Sw as decision threshold options. Structural prediction of CrsL and ApeB from *M. smegmatis* and predictions of their heterodimers with CarD or RNAP were computed using AlphaFold2 and the extension AlphaFold Multimer (72,73).

### NGS data processing and analysis

#### ChIP-seq

ChIP-seq peak calling and gene assignment was done as described previously (44). Briefly, the reads were mapped to *Mycobacterium smegmatis* genome (NCBI RefSeq NC_008596.1) with HISAT2 (74) and peaks were called by MACS2 (75). The Venn diagrams for overlapping peaks were created with BEDTools intersect (76) and matplotlib-venn package (77). The 3-way Venn diagram was created with Intervene (78). The coverage profiles of the 1 kb region around ORF start were created using deepTools (79) (computeMatrix, plotProfile) with programming libraries rtracklayer (80), Pandas (81), Matplotlib (82) and Seaborn (83).

#### RNA-seq

Read quality was checked using FastQC version 0.11.9 (https://www.bioinformatics.babraham.ac.uk/projects/fastqc/). When needed, adapters and low-quality sequences were removed using Trimmomatic 0.39 (84). Reads were aligned to the reference genome using HISAT2 2.2.1 (85) and SAMtools 1.13 (86,87). Read coverage tracks were computed using deepTools 3.5.1 (88). The DESeq2 R package (89) was used to identify differentially expressed genes at FDR ≤ 0.05.

## RESULTS

### ApeB binds to CarD in *M. smegmatis*

We started our search for CarD interacting partners using an *M. smegmatis* strain with an additional copy of FLAG-tagged CarD under an anhydrotetracycline (ATc) inducible promoter (46). Immunoprecipitation of CarD-FLAG from exponential and stationary phases revealed CarD association with RNAP subunits, σ^A^, RbpA and, notably, ApeB (MSMEG_5828), a putative protease homologous to *M. tuberculosis* PepC (90,91) (**Figure 1A**). ApeB was bound to CarD-FLAG in stationary phase (**Figure 1A, lane 4**).

**Figure 1.**
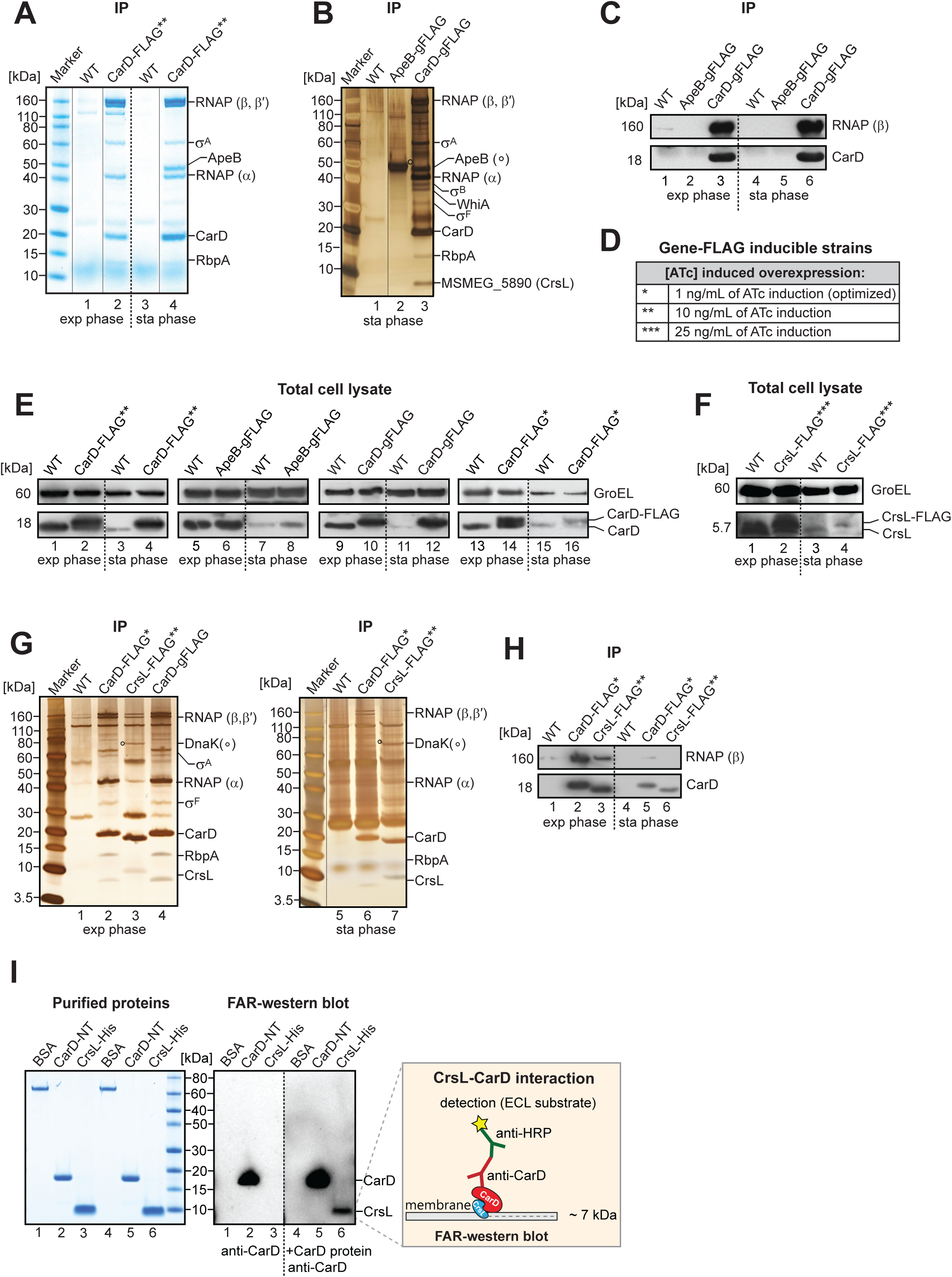
**A.** Proteins co-immunoprecipitated with CarD-FLAG from exponential and stationary phases were resolved by SDS PAGE and visualized by Coomassie. **B.** Proteins co-immunoprecipitated with ApeB-gFLAG and CarD-gFLAG from stationary phase were resolved by SDS PAGE and visualized by silver-staining. The identity of protein bands was confirmed by MALDI-FTICR mass spectrometry. The amounts of co-immunoprecipitated RNAP and CarD were determined by western blotting with anti-RNAP (β) and anti-CarD antibodies (**C**). **D.** Asterisks indicate the different ATc concentrations (ng/mL) used in the inducible FLAG-tagged strains. **E.** Western blot of total cell lysates from *M. smegmatis* wild type (WT), CarD-FLAG** (10 ng/mL ATc), ApeB-gFLAG, CarD-gFLAG and CarD-FLAG* (1 ng/mL ATc) strains from exponential and stationary phase. CarD levels were detected with anti-CarD antibody, and GroEL, as a loading control, was detected with anti-GroEL. **F.** Western blot of total cell lysates from wt (WT) and CrsL-FLAG*** (25 ng/mL ATc) from exponential and stationary phase. The levels of CrsL were detected by anti-CrsL antibody. GroEL was used as loading control and detected with anti-GroEL. **G.** Co-immunoprecipitated proteins from CarD-FLAG* (1 ng/mL ATc), and CrsL-FLAG** (10 ng/mL ATc) strains from exponential and stationary phases were resolved by SDS PAGE and visualized by silver staining. The identity of protein bands was confirmed by MALDI-FTICR mass spectrometry. **H.** The immunoprecipitated proteins from CarD-FLAG* and CrsL-FLAG** strains were detected by western blotting with anti-RNAP (β) and anti-CarD antibodies. **I.** The direct interaction of CrsL with CarD was detected *in vitro* by FAR western blotting using CarD-NT and CrsL-His purified proteins and anti-CarD antibody. The purified proteins were resolved on SDS-PAGE and stained by Coomassie. BSA and a membrane incubated without CarD protein were used as negative controls.

To confirm the CarD-ApeB interaction, we generated strains with the FLAG-tag sequences inserted into the *M. smegmatis* genome at the *carD* (CarD-gFLAG) or *apeB* (ApeB-gFLAG) loci. CarD-gFLAG immunoprecipitation confirmed the interaction with ApeB and other known interaction partners of CarD: σ^B^ and WhiA (11,92–95) (**Figure 1B, lane 3**). The reciprocal immunoprecipitation with ApeB-gFLAG yielded no CarD (**Figure 1B, 1C**).

To unravel this discrepancy, we compared the levels of CarD in the CarD-gFLAG strain, where endogenous CarD was FLAG-tagged, to those in the strain where the expression of CarD-FLAG was induced by different ATc concentrations which resulted in altered CarD-FLAG level (**Figure 1D**), and wild type (wt) strain. In wt and CarD-FLAG* strain (where the lowest level of ATc, 1 ng/mL, was used to induce CarD-FLAG), as well as ApeB-gFLAG strain, the CarD level dramatically decreased between exponential and stationary phases (**Figure 1E**, compare *e. g.* **lane 1 *vs.* lane 3**). In contrast, in strains with CarD-FLAG** (where the higher level of ATc, 10 ng/mL, was used for CarD-FLAG induction) and CarD-gFLAG, the level of CarD remained high also in stationary phase (**Figure 1E**, **lanes 4 *and* 12**). This raised the question of why CarD level increased in stationary phase in the CarD-gFLAG strain, in which CarD is expressed from its endogenous promoter. We noticed an antisense RNA expressed from the *ispD-carD* locus in our RNA-seq data (41). This antisense RNA of *carD* (*AscarD* RNA) was recently shown to be expressed in stationary phase and to negatively regulate CarD expression (38). In the CarD-gFLAG constructed strain, the addition of the sequence encoding FLAG-tag to the *carD* genome locus disrupted expression of *AscarD* RNA, which resulted in an increased level of the CarD protein (**Figure 1E, lane 12** and **Supplementary Figure 1A**).

Therefore, the high level of CarD-FLAG was responsible for the observed interaction between ApeB and CarD. The ApeB-CarD interaction in stationary phase is detected only when CarD is overexpressed. To conclude, ApeB binds CarD only when the CarD level is elevated. While further investigation is needed to determine the conditions under which ApeB binds CarD in the wild type strain, the immunoprecipitation experiments revealed another promising candidate protein associated with CarD: MSMEG_5890 (**Figure 1B**).

### CrsL binds to CarD in *M. smegmatis*

MSMEG_5890 encodes a small protein (predicted Mw 5.7 kDa), which we named “CrsL” (<u>C</u>arD <u>R</u>NA polymerase <u>s</u>mall linker) (**Figure 1B**). The function of CrsL is unknown. Unlike ApeB, CrsL interacted with CarD-FLAG not only when the CarD-FLAG levels were elevated (**Figure 1B**), but also when CarD-FLAG levels were optimized to be comparable to the endogenous CarD level in wt strain (**Figure 1E, lanes 14 *and* 16, Figure 1G, lanes 2 *and* 6**). Under these conditions, CrsL peptides were highly abundant in CarD-FLAG* immunoprecipitates in both exponential and stationary phases as detected by MALDI-FTICR mass spectrometry (**Supplementary Table 1**).

To validate the CrsL-CarD interaction, we constructed a strain with FLAG-tagged CrsL under an ATc inducible promoter (46). We generated our own mouse anti-CrsL antibody and verified the CrsL-FLAG expression. Even with induction of 25 ng/mL ATc (indicated by *** in **Figure 1F**), CrsL-FLAG levels were comparable to endogenous CrsL in wt (**Supplementary Figure 1B, lane 1 *vs.* 2**). CrsL-FLAG** immunoprecipitated CarD from both exponential and stationary phase cells as detected by mass spectrometry (**Figure 1G, lanes 3 *and* 7**), and by western blotting with the anti-CarD antibody (**Figure 1H**). Additionally, in exponential but not in stationary phase, RNAP β, β*’* and α subunits co-immunoprecipitated with CrsL-FLAG** (**Figure 1G, lane 3 *and* 7** and **Figure 1H, lane 3 *and* 6**). In both phases, CrsL interacted with the DnaK chaperon protein (MSMEG_0709) (**Figure 1G, lanes 3 *and* 7**).

Finally, to determine whether the CrsL-CarD interaction is direct or mediated by another protein, we performed FAR-western blotting (51) using *in vitro* purified CarD and CrsL proteins. With the anti-CarD antibody, we detected CarD associated with the nitrocellulose membrane-bound CrsL (**Figure 1I, lane 6**), indicating that CrsL directly binds to CarD.

### The majority of CrsL co-sediments with CarD

In exponential phase, CrsL and RNAP were co-immunoprecipitated with CarD-FLAG. Reciprocally, CarD and RNAP were co-immunoprecipitated with CrsL-FLAG (**Figure 1G**, **1H**). To determine whether CrsL primarily associates with RNAP or with CarD, we performed glycerol gradient ultracentrifugation using exponential and stationary phase *M. smegmatis* lysates. The gradient was then fractionated and the distribution of CrsL, RNAP, σ^A^, CarD, and RbpA analyzed by western blotting.

In exponential phase, CrsL mainly sedimented in the top fractions (1–5) where CarD was also predominantly found (**Supplementary Figure 1E**). Fractions 4 and 5 contained RNAP (**Supplementary Figure 1E**), but the majority of CrsL co-sedimented with CarD in fractions 1-3. Only a subset of CrsL molecules co-sedimented with both CarD and RNAP in fractions 4 and 5, indicating that the majority of CrsL is not bound to RNAP.

In stationary phase, CrsL was not detected in the top fractions where most of CarD sedimented. CrsL sedimented in the bottom fraction together with a minor amount of CarD (fraction 20) (**Supplementary Figure 1E**). Although *M. tuberculosis* RNAP can oligomerize *in vitro* into supramolecular complexes (96) and RNAP was partially detected in the gradient’s bottom fraction (**Supplementary Figure 1E**), RNAP was not co-immunoprecipitated with CrsL-FLAG in stationary phase (**Figure 1G**). This suggests that CrsL likely participates in other complexes that do not include RNAP, possibly involving the DnaK chaperone, a major CrsL-interacting protein (along with CarD) in stationary phase (**Figure 1G, lane 7**).

### CrsL is evolutionarily conserved in actinobacteria

To further characterize CrsL, we explored the evolutionary relationship between its homologs across diverse bacterial species. We conducted a multiple sequence alignment with selected CrsL homologs, identifying the 20-45 aa residue region as the most conserved part of CrsL (**Figure 2A**, **2B** and **Supplementary Table 2**). CrsL is conserved in many actinobacterial species including *Mycobacteria*, *Nocardia, Streptomyces, Corynebacteria* and *Rhodococcus* (**Figure 2A** and **Supplementary Table 3**). CrsL homologs are found in species such as *M. tuberculosis* (*Rv3489*), *M. bovis* (*Mb3519*) and *M. marinum* (*MMAR_4977*) where they are annotated as hypothetical proteins with unknown function (97,98).

**Figure 2.**
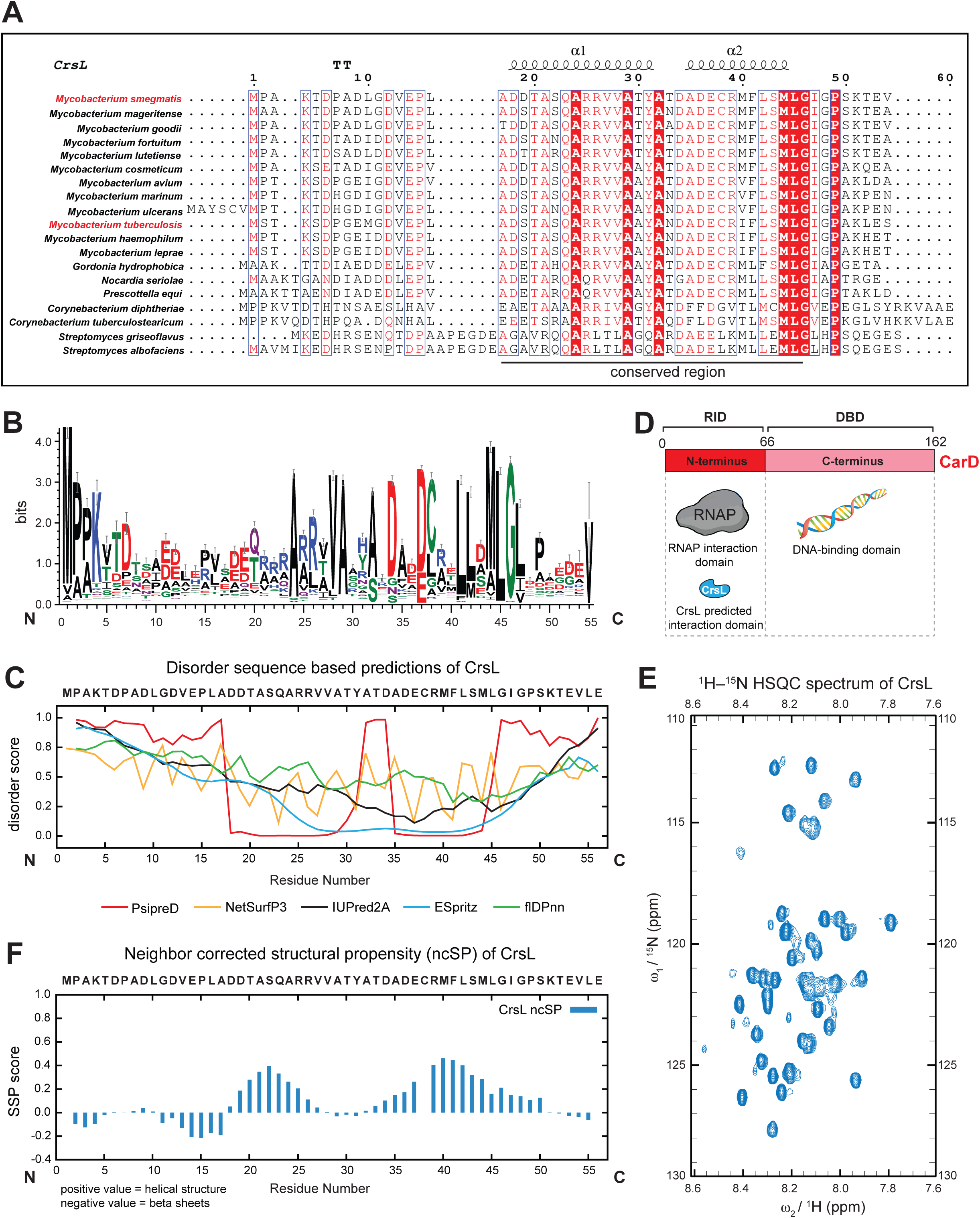
**A.** Multiple sequence alignment of selected CrsL homologs in actinobacteria. The conserved regions and the secondary structure elements are depicted above the alignment. The protein names are listed in **Supplementary Table 2**. **B.** Multiple Sequence Alignment of CrsL sequence by DeepMSA2 showing the highly conserved region (20-45 residues) within 770 entries of homologous bacterial sequences. **C.** Disorder sequence-based prediction of CrsL using different web-tools. Tools’ names are indicated below the figure. The values above 0.4 are considered as intrinsically disordered regions. CrsL protein does not have an ordered structure with exceptions of PsipreD and ESpritz predictions. **D.** A scheme showing the domains of CarD proteins where RNAP interacts with the N-terminal domain, which is the same interaction domain predicted for CrsL binding. **E.** The 2D ^1^H–^15^N hetero-nuclear single quantum coherence (HSQC) spectrum of CrsL. The proton resonances have a narrow distribution pointing towards disordered state of the protein. **F.** Neighbor Corrected Structural Propensity (ncSP) of CrsL calculated from the chemical shifts of the protein backbone. Relatively low values point to a disordered state with a slight propensity towards 2 alpha-helical regions. The cysteine residues (as Cys38 in CrsL) are not computed by the ncSP default settings.

The CrsL encoding gene is organized in the *MSMEG_5890-MSMEG_5892* two-gene operon and is transcribed from a single promoter located upstream *crsL* (99). This organization is highly conserved in other mycobacterial species including *M. tuberculosis* (100) (**Supplementary Figure 2**). The second gene of the operon, *MSMEG_5892* (*otsA*), which is not essential in *M. smegmatis* (101,102), encodes α,α-trehalose-phosphate synthase. Trehalose is a disaccharide important for the biosynthesis of the mycobacterial cell wall (103–105).

### CrsL is an intrinsically disordered protein

To probe the secondary structures of CrsL and obtain insights into its potential three-dimensional arrangement, we first used protein structure predictions. The combined results from several prediction tools, namely PsipreD (67), NetSurfP3 (68), IUPred2A (69), ESpritz (70) and flDPnn (71), showed low values for secondary structures. Three prediction tools (NetSurfP3, flDPnn, IUPred2A) consistently assigned ambiguous values to most residues. On the contrary, PsipreD and ESpritz predicted two conserved regions of CrsL (aa 20-27 and 34-48) to be well-ordered (**Figure 2C**). Similarly, AlphaFold3 (72) predicted these two regions to be structured (**Supplementary Figure 3A and 3E, in green**). This suggests formation of ordered structures in complexes with interacting partner(s). The CrsL-CarD interaction predicted by AlphaFold Multimer (72,73) showed that the ordered part lies closer to the C-terminus of CrsL and the N-terminal domain of CarD (RNA interaction domain, RID) (**Figure 2D** and **Supplementary Figure 3A-D**). The predicted interaction site is close to the hydrophobic pocket of CarD’s N-terminal domain and the interaction surface is mediated mostly by aa residues 1-60 with the main hydrophobic residues identified as Tyr55, Val48 and Leu44 (**Supplementary Figure 3D**).

Next, we purified CrsL labeled with ^13^C and ^15^N isotopes and measured its NMR spectra and performed backbone assignment. We successfully identified all of the peaks corresponding to all atoms that form the CrsL backbone. The 2D ^1^H–^15^N hetero-nuclear single quantum coherence (HSQC) spectra were used to monitor the individual amide-proton signals of the protein (**Figure 2E**). Based on the obtained spectrum, the proton resonances exhibit a narrow distribution typical of many intrinsically disordered proteins (**Figure 2C**), including residues forming the two regions predicted as structured, Furthermore, we computed the neighbor corrected structural propensity (ncSP) and secondary structure propensities (SSP) to obtain more detailed structural information. The low ncSP/SSP values indicate that CrsL is an intrinsically disordered protein when free in solution (**Figure 2F** and **Supplementary 3E, in red**).

### CrsL is a potential mycobacterial transcription factor

As CrsL was found in complex with RNAP and CarD in exponential phase (**Figure 1G**), we employed chromatin immunoprecipitation sequencing (ChIP-seq) using CrsL-FLAG to determine if CrsL associates with the bacterial chromosome. Additionally, we compared the genomic binding sites of CrsL with that of CarD, RbpA, RNAP, and σ^A^/σ^B^ (44,45) under the same experimental conditions. All ChIP-seq data and detected peaks are available at the msmegseq.elixir-czech.cz webpage with the integrated IGV genome browser (106).

The results showed that CrsL indeed associates with the genome. We detected comparable numbers of CrsL peaks in the *M. smegmatis* genome relative to CarD peaks (315 vs 406, **Figure 3A** and **Supplementary Table 4**). CrsL and CarD peaks were highly enriched at the 5 ends of genes − regions that correspond to gene promoters. An example of representative ChIP-seq data is shown in **Figure 3B**.

**Figure 3.**
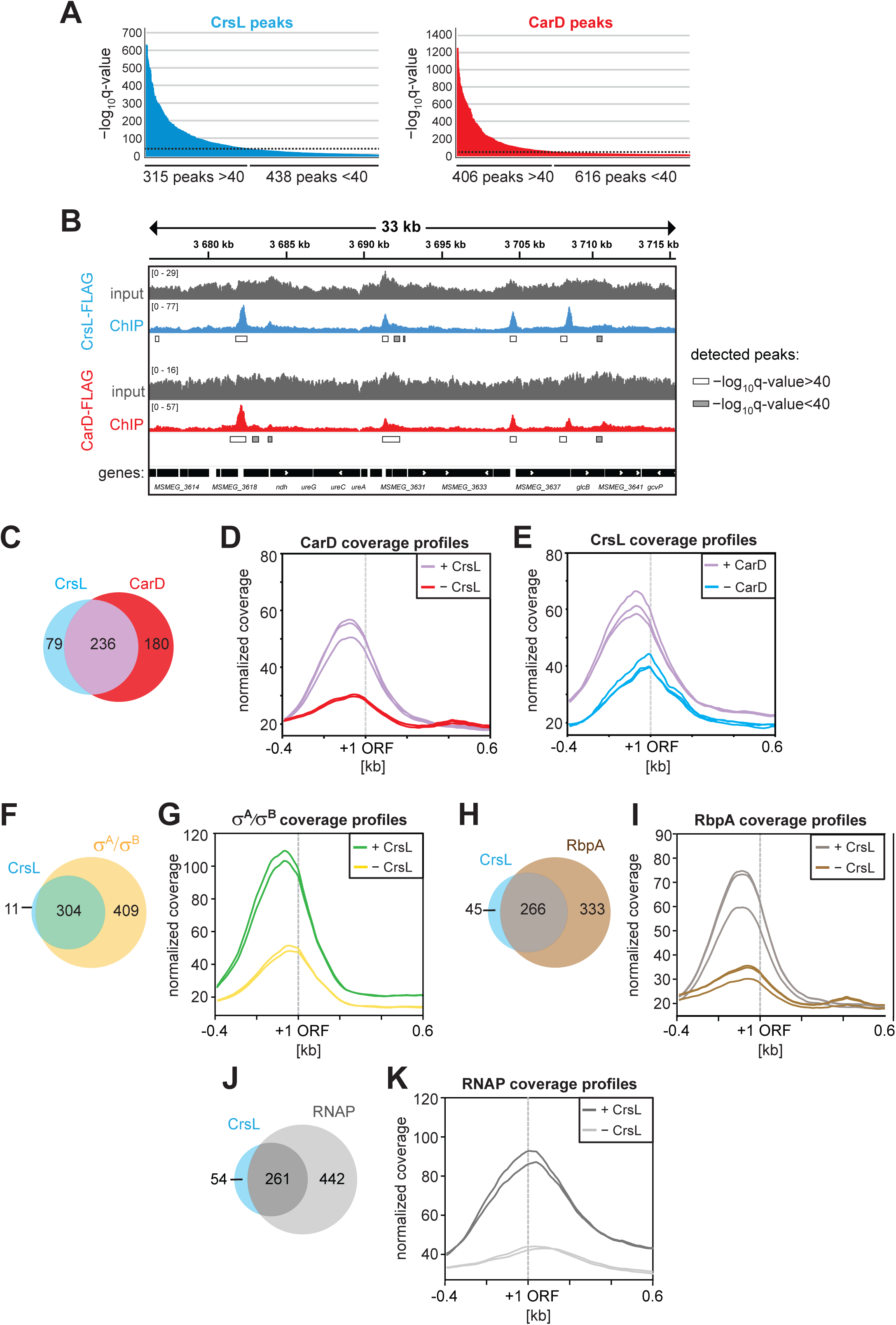
**A.** ChIP-seq peaks detected with CrsL-FLAG and CarD-FLAG. The experiments were done in biological triplicates. The peaks are divided according to significancy cutoff set as 40 and the higher −log_10_ q-value value indicates higher statistically significant peaks. **B.** An example ChIP-seq detected peaks for CrsL-FLAG versus CarD-FLAG within 33 kb region. The significancy of the peaks are indicated by boxes (white/grey) below the detected peaks. **C.** Venn diagram showing overlap of CrsL and CarD binding sites derived from ChIP-seq peaks. **D.** ChIP-seq coverage profiles of CarD for genes either associated with both CarD and CrsL (+CrsL) or genes associated only with CarD (−CrsL) in 1 kb region around ORF for individual samples (lines). Genes significantly associated with both CarD and CrsL at their promoters tend to have higher CarD peak coverage than those lacking CrsL association (purple lines). **E.** ChIP-seq coverage profiles of CrsL for genes either associated with both CrsL and CarD (+CarD) or genes associated only with CrsL (−CarD) in 1 kb region around ORF for individual samples (lines). Genes significantly associated with both CrsL and CarD at their promoters tend to have higher CarD peak coverage than those lacking CrsL association (purple lines). **F.** Venn diagram showing overlap of CrsL and σ^A^/σ^B^ binding sites derived from ChIP-seq peaks. **G.** ChIP-seq coverage profiles of σ^A^/σ^B^ for genes either associated with both σ^A^/σ^B^ and CrsL (+CrsL) or genes associated only with σ^A^/σ^B^ (−CrsL) in 1 kb region around ORF for individual samples (lines). **H.** Venn diagram showing overlap of CrsL and RbpA binding sites derived from ChIP-seq peaks. **I.** ChIP-seq coverage profiles of RbpA for genes either associated with both RbpA and CrsL (+CrsL) or genes associated only with RbpA (−CrsL) in 1 kb region around ORF for individual samples (lines). **J.** Venn diagram showing overlap of CrsL and RNAP binding sites derived from ChIP-seq peaks. **K.** ChIP-seq coverage profiles of RNAP for genes either associated with both RNAP and CrsL (+CrsL) or genes associated only with RNAP (−CrsL) in 1 kb region around ORF for individual samples (lines).

Around 75 % of the significant peaks (−log_10_ q-value > 40) detected for CrsL overlapped with CarD peaks (**Figure 3C**). Especially promoter regions with high CarD occupancy were almost always associated with CrsL (**Figure 3D**). Likewise, promoter regions with high CrsL occupancy were associated with CarD (**Figure 3E**). This positive correlation between CrsL and CarD suggests that the presence of one protein enhances the binding of the other protein to the promoters. Nevertheless, both CrsL and CarD can also interact with promoters in the absence of each other (**Figure 3C, blue *vs.* red colors**).

In the ChIP-seq data, only 79 of CrsL-associated genomic loci were not bound by CarD (**Figure 3C**). However, we noticed that 56 of these 79 CrsL peaks actually overlap with low-significant CarD (−log_10_ q-value < 40) peaks that were originally excluded from the analysis. The remaining subset of 23 CrsL peaks do not overlap with any detectable CarD peaks but exclusively overlap with σ^A^/σ^B^ peaks (no RNAP, CarD, and RbpA peaks). An example is the promoter of the genes *MSMEG_1821* (encoding acyl-CoA dehydrogenase) or *MSMEG_5136* (annotated as helix-turn-helix motif protein) (**Supplementary Figure 4A**).

The majority of CrsL-bound genomic loci were also associated with σ^A^/σ^B^, RbpA and RNAP (**Figure 3F**, **3H** and **3J**, respectively). Moreover, promoters associated with CrsL displayed higher coverage of σ^A^/σ^B^, RbpA and RNAP peaks than promoters without CrsL (**Figure 3G**, **3I** and **3K**, respectively). Interestingly, across the entire *M. smegmatis* genome, we identified only 11 peaks of CrsL that did not overlap with σ^A^/σ^B^ (**Figure 3F**). This suggests that CrsL predominantly associates with promoter regions of σ^A^/σ^B^-dependent genes (**Figure 3G**).

The CrsL peaks, similar to CarD peaks, were detected at the promoters of genes involved in transcription regulation, including *carD*, *rpoB*, and *sigA* genes (**Figure 4A**). CrsL also binds to its own promoter (**Figure 4A**), rRNA promoters (**Supplementary Figure 4B)**, tRNA genes and ribosomal protein coding genes (**Supplementary Table 4**). Based on Gene Ontology Term analysis (107), many CrsL-bound genes are involved in *Protein biosynthesis*, *Amino-acid biosynthesis*, *Tricarboxylic acid cycle* biological processes. We generated average ChIP-seq profiles of CrsL and CarD for 200 highly expressed genes as well as for 200 genes with low to no expression in exponential phase (41) (**Figure 4B**, **4C**, respectively; for expression profiles of the two gene groups see **Supplementary Figure 4C**). CrsL was associated with highly expressed genes and was notably absent from non-transcribed or lowly expressed genes, as was CarD (**Figure 4B** and **4C**). Globally, CrsL interacted with promoters of actively transcribed genes similarly to CarD, RNAP or σ^A^/σ^B^ (**Figure 4D**) and majority of CrsL-associated promoters interacted also with CarD, RNAP, RbpA and σ^A^/σ^B^ (**Figure 4E**, **4F**). These data suggest that CrsL is a transcription factor that binds to promoters of highly expressed genes in *M. smegmatis*.

**Figure 4.**
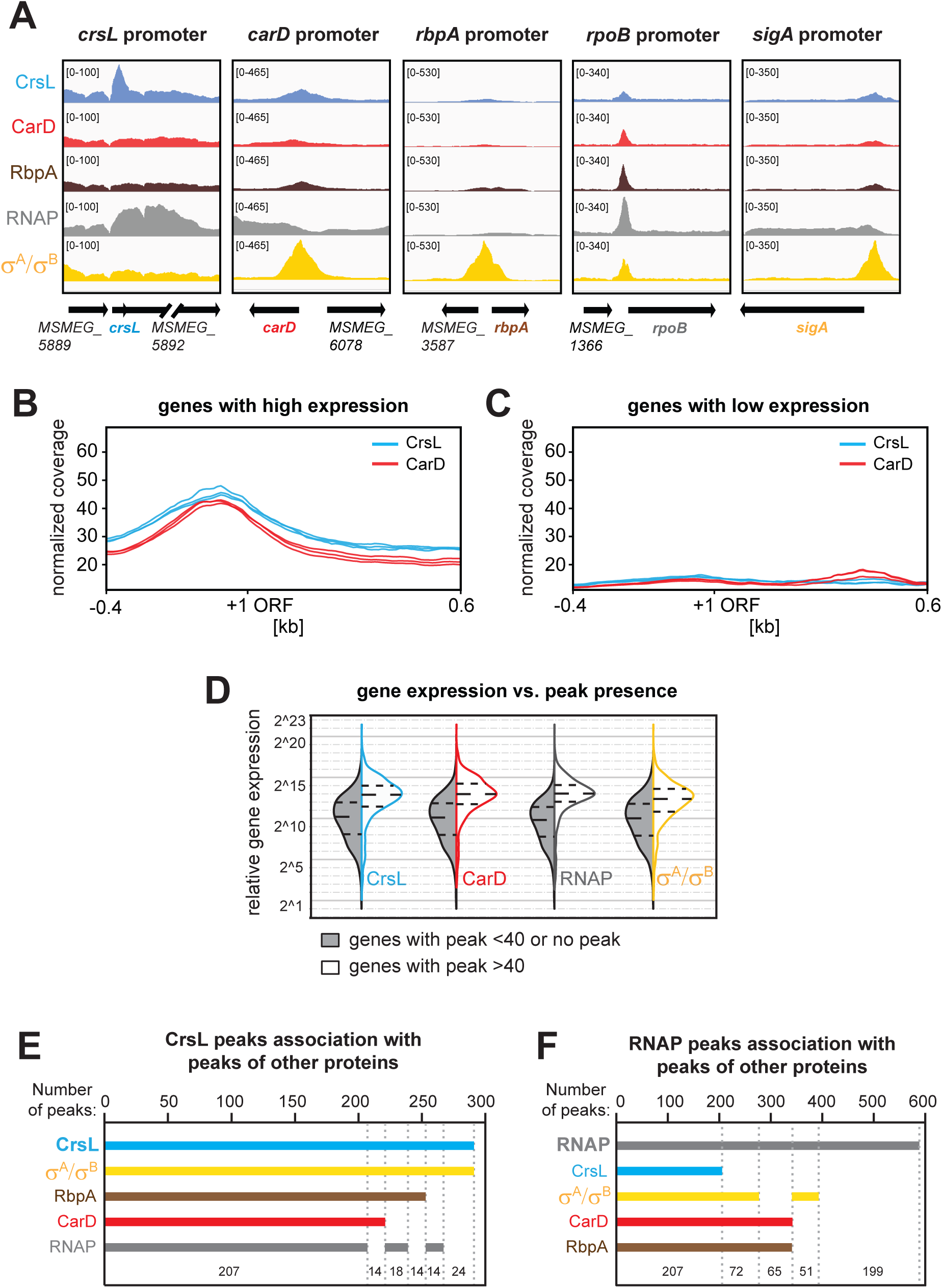
**A.** ChIP-seq detected peaks for CrsL, CarD, RbpA, RNAP and σ^A^/σ^B^ on their promoters. For comparison, peaks height for all datasets were set to the same value based on the highest peak score. Average of CrsL (in blue) and CarD (in red) peak profiles (each line represents one biological replicate) over 200 genes with highest expression (**B**) and 200 genes with no or lowest expression (**C**) in the exponential phase in *M. smegmatis* (49). 1000 bp region is shown with +1 representing start site of ORF and -0.4 kb upstream and +0.6 kb downstream of ORF. **D.** Violin plots showing relative gene expression level for genes with/without significant CrsL, CarD, RNAP and σ^A^/σ^B^ association. The white color (right sides) of violin plots shows expression of genes with peaks −log_10_ q-value >40 while the grey color (left sides) of violin plots shows expression of genes with no or less significant peaks (−log_10_ q-value <40). Dashed lines show lower and upper quartiles. **E.** Plot showing intersections of CrsL peaks with peaks of σ^A^/σ^B^ or RbpA or CarD or RNAP. CrsL predominantly associates with majority of gene promoters with a peak profile intersects with CarD, RbpA, RNAP and more with σ^A^/σ^B^. **F.** Plot showing intersections of RNAP peaks with peaks of CrsL or σ^A^/σ^B^ or CarD or RbpA. The 199 RNAP peaks that are not present in CrsL, RbpA, CarD and σ^A^/σ^B^ likely represent those peaks of RNAP located within gene regions or at the 3’ ends of genes. The major set of RNAP peaks (n=207) overlaps with peaks of CarD, RbpA, σ^A^/σ^B^ and CrsL.

### CrsL protein cannot be efficiently depleted in *M. smegmatis*

We generated strains containing an ATc-inducible CRISPR system (48) to knockdown CrsL, CarD, or σ^A^ (the latter two essential proteins (9,33) were used as positive controls). Upon ATc addition, the mRNA levels of *crsL* and *carD* genes were efficiently depleted (>90 % depletion) in both exponential and stationary phases as confirmed by RT-qPCR (**Figure 5B**). We note, however, that at the protein level, CrsL was depleted by 70 % and 50 % in exponential and stationary phase, respectively, whereas depletion of CarD was more efficient (>90 % depletion) (**Figure 5C**, **5D**). Depletion of CarD or σ^A^ resulted in growth inhibition zones for the respective strains, while no inhibition zone was observed with the CrsL depletion strain (**Figure 5A**). Therefore, the reducing CrsL protein level by half did not affect *M. smegmatis* growth.

**Figure 5.**
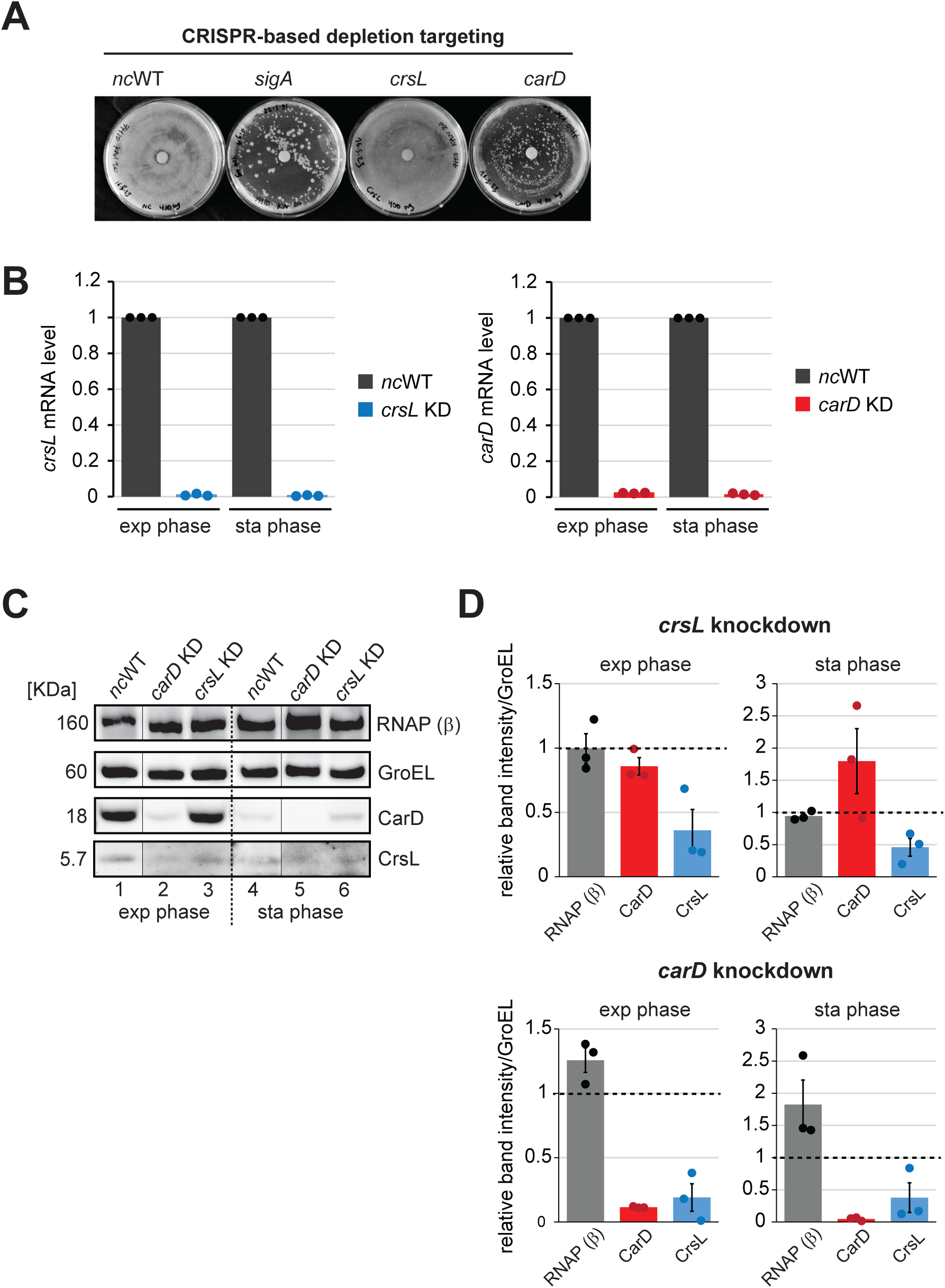
**A.** Depletion of *crsL* using CRISPR. The depletion was induced by adding 4 µL of 100 ng/mL ATc onto discs placed on plates streaked with the bacterial strains. **B.** The mRNA levels of depleted *crsL* and *carD* genes in *crsL* knockdown and *carD* knockdown strains, respectively, in exponential and stationary phase were measured by RT-qPCR with gene specific primers. The data was normalized to spike-in control and negative control (*nc*WT) was set as 1. Three-four biological replicates were used, and error bars represent SEM. **C.** The protein levels of RNAP (β), CarD and CrsL upon depletion of *crsL* and *carD* detected by western blotting using anti-RNAP (β), anti-CarD and anti-CrsL antibodies. GroEL level was used as loading control and *nc*WT was used as a negative control. **D.** The quantified western blot bands of protein levels of RNAP (β), CarD and CrsL upon depletion of *crsL* and *carD*, from biological triplicates. The data was normalized to GroEL level, and the negative control (*nc*WT) was set as 1. Error bars represent SEM.

### CrsL level is decreased in stationary phase and is dependent on CarD

Both CarD and CrsL protein levels decreased in stationary phase (**Figure 1F**). When the CarD protein was depleted, the CrsL protein level was reduced (**Figure 5C**, **5D**). Conversely, when the CarD level was artificially upregulated in the CarD-gFLAG strain during stationary phase, the CrsL protein level increased (**Supplementary Figure 1C, lane 8**). Therefore, the amount of CrsL depends on the amount of CarD. Consistent with this, more CrsL was co-immunoprecipitated with CarD-FLAG in exponential phase; where CarD levels are higher than in stationary phase (**Figure 1G, lane 2 and 4 *vs.* lane 6**). Additionally, more CrsL was co-immunoprecipitated when CarD level was increased in CarD-gFLAG strain during stationary phase (**Supplementary Figure 1D, lane 8**). The higher CrsL levels also co-immunoprecipitated more CarD in exponential phase (**Figure 1G, lane 3 *vs.* lane 7** and **Supplementary Figure 1D, lane 3 *vs.* lane 7**). Thus, the formation of the CarD-CrsL complex is mainly regulated by the levels of the interacting proteins during cell growth.

### Identification of CrsL- and CarD-regulated genes in *M. smegmatis*

To determine the effects of *crsL* depletion on the *M. smegmatis* transcriptome, we performed RNA-seq using the *crsL* knockdown strain from exponential and stationary phase. In parallel, we performed RNA-seq with the *carD* knockdown strain to compare the differentially expressed genes (DEGs) regulated by CrsL and CarD.

After CrsL depletion in exponential phase, relatively few genes were affected (|Log_2_FC| ≥ 0.5, FDR-corrected *p*-value ≤ 0.05): 43 genes significantly increased expression; while 90 genes decreased expression **(Figure 6A**). In stationary phase, 141 genes significantly increased expression while 320 genes decreased their expression (**Figure 6B**). In either phase, CrsL depletion did not affect the *carD* mRNA level. The relatively small number of affected genes could be due to the less efficient depletion of the CrsL protein. The differentially expressed genes likely represent the tip of the iceberg – genes requiring CrsL the most for their regulation.

**Figure 6.**
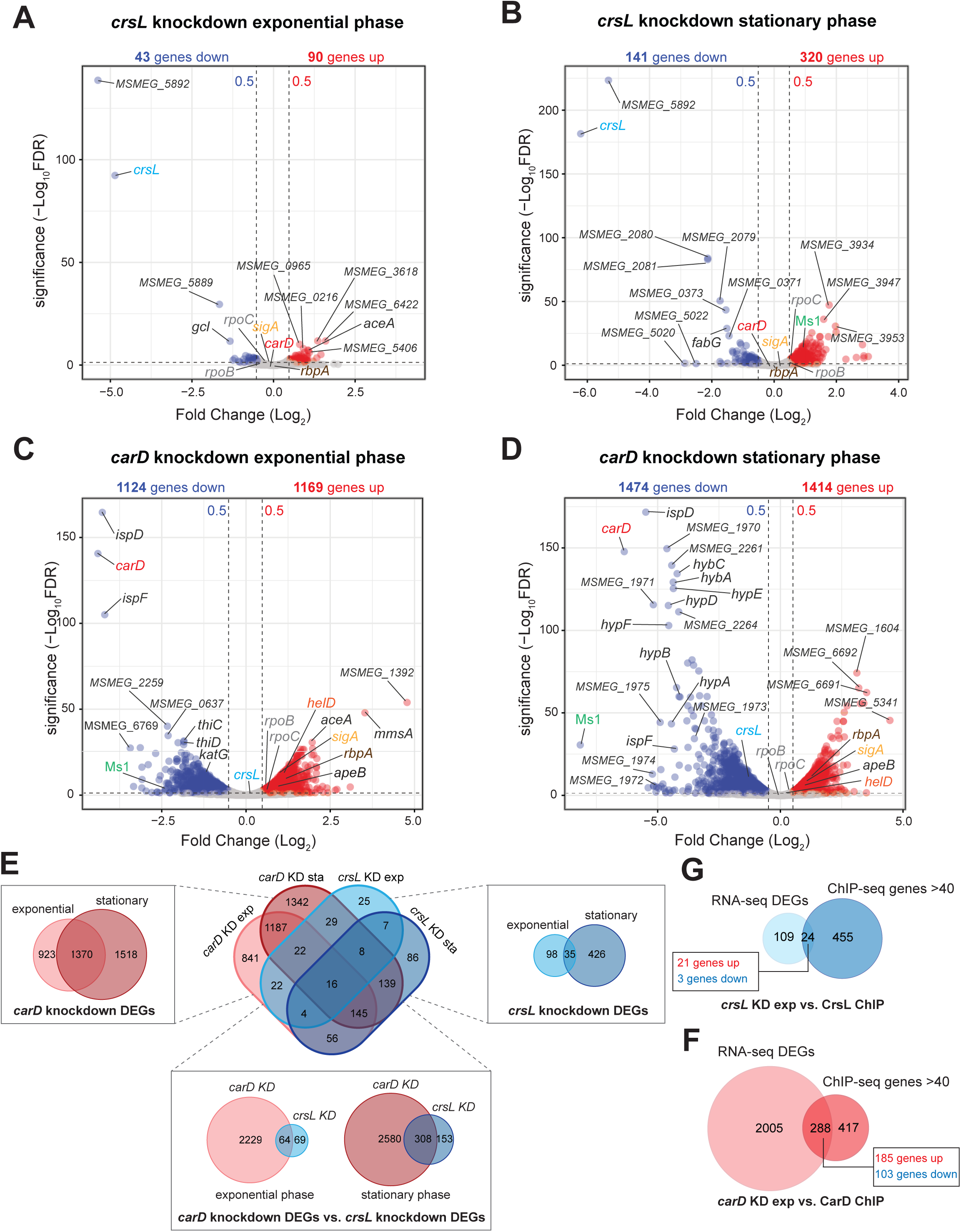
Volcano plots showing fold change (FC, log_2_) against significance (−log_10_FDR corrected *p*-value) of transcripts identified by RNA-seq analysis of *crsL* knockdown or *carD* knockdown in exponential phase (**A, C**; respectively) and *crsL* knockdown or *carD* knockdown in stationary phase (**B, D**; respectively). Blue dots indicate significant genes with a FC ≤ −0.5; red dots indicate significant genes with a FC ≥ 0.5; grey dots indicate unchanged genes. The significancy cutoff (−log _10_FDR corrected *p*-value) was set to 1.3 (i.e. FDR ≤ 0.05). **E.** Venn diagrams showing multiple overlap of differentially expressed genes (DEGs) between different datasets of *carD* knockdown and *crsL* knockdown. **F.** Venn diagrams showing overlap of *crsL* knockdown exponential phase DEGs with CrsL ChIP-seq associated genes. **G.** Venn diagrams showing overlap of *carD* knockdown exponential phase DEGs with CarD ChIP-seq associated genes.

To the contrary, after CarD depletion, nearly one third of the genes in the genome changed expression: 1,124 genes significantly increased expression while 1,169 genes decreased expression in exponential phase (**Figure 6C, Supplementary Table 5**). In stationary phase, 1,474 genes significantly increased expression while 1,414 genes, including *crsL*, decreased expression (**Figure 6D**). Interestingly, the most affected gene in stationary phase was Ms1, a binding partner of the RNAP core (40,41). The effects of CarD and CrsL depletions in stationary phase were unexpected as their protein levels are already highly reduced compared to exponential phase (**Figure 1E**, **1F**).

When we compared genes regulated by CrsL and CarD, 64 genes were affected by both CrsL and CarD depletion in exponential phase (∼50 % of *crsL* DEGs, **Figure 6E**). In stationary phase, 308 genes were affected by both CrsL and CarD depletion (∼66 % of *crsL* DEGs, **Figure 6E**).

Most of the differentially expressed genes are indirectly regulated by CrsL and CarD in the exponential phase. ChIP-seq detected CrsL association with 24 out of the 133 genes that changed expression upon CrsL depletion (**Figure 6F, Supplementary Table 6**). A similar scenario was observed for CarD - 288 genes (out of 2,293 genes that changed expression upon CarD depletion) were associated with a CarD (**Figure 6G, Supplementary Table 6**). Additionally, many promoters were occupied by CrsL or CarD, but the expression of the genes under these promoters was not altered after CrsL or CarD depletion, respectively (**Figure 6F**, **6G**).

Interestingly, CrsL directly regulated *MSMEG_5773* (encoding DesA, fatty acid desaturase) and *MSMEG_1930* (encoding DEAD/DEAH box RNA helicase). The DesA enzyme introduces double bonds into fatty acid chains, producing unsaturated fatty acids. This process is crucial for enhancing membrane fluidity at low temperature (108–113). DEAD/DEAH RNA box helicase remodels the RNA secondary structures formed during cold stress (114–119). CrsL acts as a repressor for these genes; it associates with their promoters, and expression of both genes increased following CrsL depletion. By repressing these two genes, CrsL may support growth at elevated temperatures.

### CrsL improves *M. smegmatis* growth at elevated temperature

To test the ability of CrsL to enhance the growth of *M. smegmatis* under increased temperature, we measured the growth curves of CrsL-depleted cells (*crsL* knockdown) and the control (*nc*WT) strain at 37°C and 45°C (heats stress was induced during mid-exponential phase). At 37°C, there was no significant difference between the strains (**Figure 7A**). However, at 45°C, CrsL-depleted cells showed slower growth compared to the control (**Figure 7A**). These findings indicate that CrsL functions as a transcriptional regulator that supports growth at elevated temperatures. A scheme representing the role of CrsL in modulating the cell membrane and RNA structures during temperature changes is shown in **Figure 7B**.

**Figure 7.**
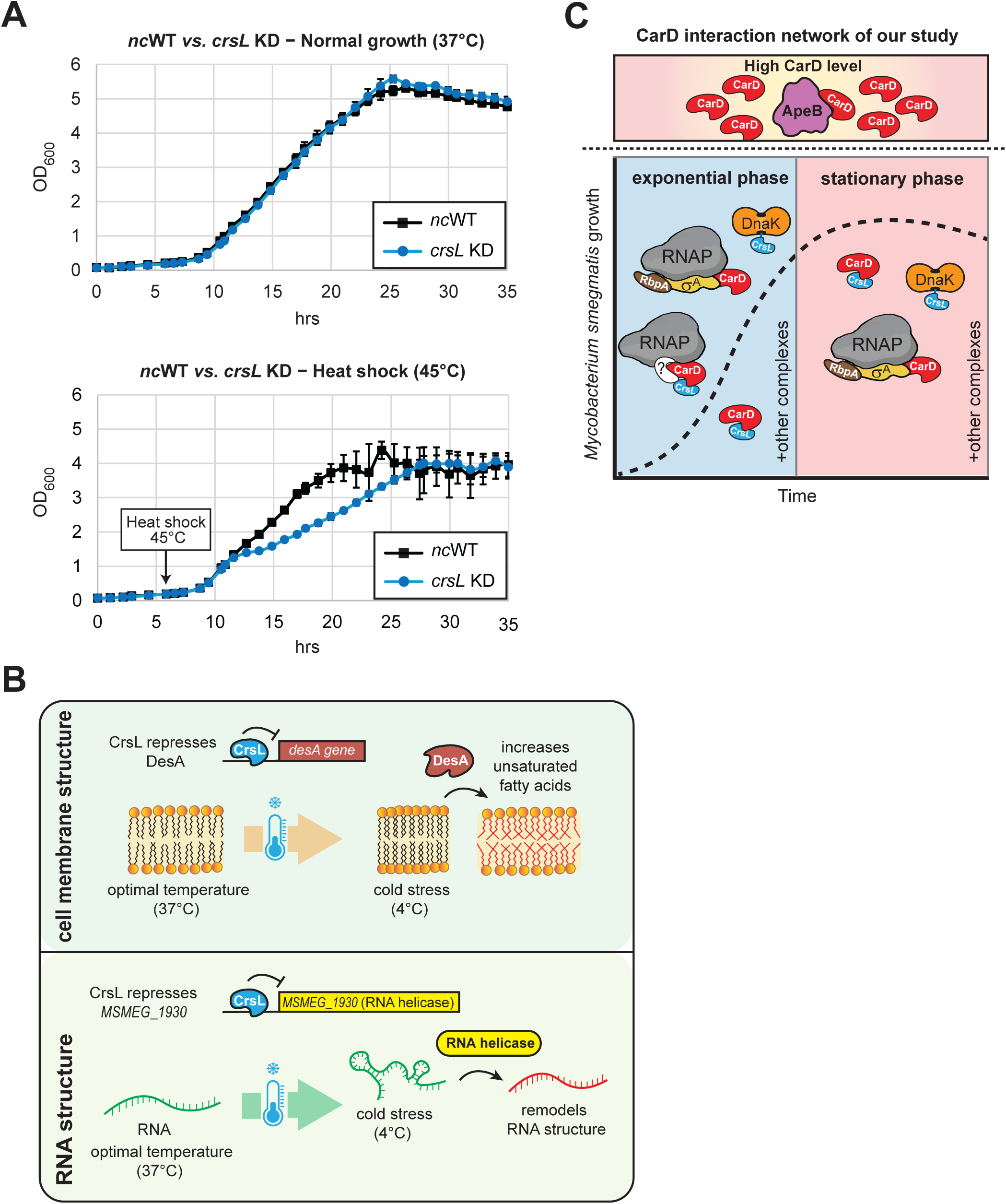
**A.** Growth curves of *crsL* depletion strain versus the negative control (*nc*WT) strain from biological triplicates incubated at 37°C (upper plot) or at 45°C (lower plot). The cells were grown in BioSan RTS-8 Multi-Channel Bioreactor and OD_600_ was measured over the growth. Error bars represent SEM. **B.** The role of CrsL in modulating the cell membrane and RNA structures by regulating the expression of DesA and DEAD-box helicase during temperature changes. **C.** A schematic representation for CarD interaction network of our study and proposed functions of CrsL.

We conclude that CrsL is a novel transcription factor that interacts with CarD and affects gene expression during both growth phases. CrsL regulon largely overlaps with that of CarD, suggesting a regulatory interplay between these two proteins. Notably, CrsL specifically represses two genes important for survival during cold stress, allowing *M. smegmatis* to grow effectively at higher temperatures.

## DISCUSSION

In this study, we expanded the interaction network of the essential mycobacterial transcription factor, CarD, by identifying two new interacting partners - ApeB and CrsL in *M. smegmatis* (**Figure 7C**).

ApeB is a putative aminopeptidase that cleaves proteins from the amino terminus. We showed that ApeB binds to CarD only when CarD levels are elevated. The expression of *apeB* homolog in *M. tuberculosis* (*pepC, Rv0800*) was significantly increased (∼2-fold) in the strain where mutated CarD had increased affinity to RNAP (35). In our RNA-seq dataset, the transcript level of *apeB* was altered upon CarD depletion both in exponential and stationary phase, even though CarD was not associated with the *apeB* promoter (**Figure 6C**, **6D**). Therefore, both the ApeB-CarD interaction and the level of ApeB itself might depend on the amount of CarD (either free or RNAP-bound) in mycobacteria. The specific function of ApeB and its homolog in *M. tuberculosis* is currently unknown, but our data suggest that this aminopeptidase is able to interact with CarD and therefore might affect transcription under conditions where CarD levels are increased. AlphaFold predicted structural interaction between ApeB and CarD is shown in **Supplementary Figure 3F**.

In contrast to ApeB, CrsL binds to the cellular levels of CarD. The CrsL-CarD interaction is direct and both proteins associate with RNAP. What is the role of CrsL in mycobacteria? CrsL is conserved among actinobacteria. In *M. tuberculosis*, *crsL* homolog (*Rv3489*) is not essential (120,121) and its expression is predicted to be regulated by IdeR (122); the iron-dependent regulator, which represses genes involved in iron uptake and maintains iron homeostasis. In *M. bovis* BCG, the expression of the *crsL* homolog (*BCG_3553*) is activated by RaaS (regulator of <u>a</u>ntimicrobial-<u>a</u>ssisted <u>s</u>urvival) transcription factor, which is important for mycobacterial long-term survival (123). CrsL may therefore play a role in adapting to environmental changes, such as fluctuating iron levels or conditions associated with long-term growth.

In *M. smegmatis*, CRISPR-mediated depletion of *crsL* mRNA was highly effective (almost 99 %), yet the reduction in CrsL protein was suboptimal, indicating that CrsL is regulated post-transcriptionally. Bacteria appears to compensate for decreased *crsL* mRNA level by enhancing its translation or stabilizing CrsL protein. CrsL protein level depends on CarD, which is also regulated post-transcriptionally by an antisense RNA of *carD* (*AscarD* RNA) and the Clp protease (38). This suggests that the bacteria actively monitor both CarD and CrsL to maintain their optimal levels necessary for gene expression.

ChIP-seq data revealed that CrsL binding sites in the *M. smegmatis* genome overlap with those of CarD as well as RbpA, σ^A^/σ^B^, and RNAP. CrsL predominantly binds to promoter regions. CrsL showed a similar binding pattern to CarD, and both proteins preferentially interacted with actively transcribed genes. Considering the interaction of CrsL with CarD and their similar expression profiles during *M. smegmatis* growth (**Figure 1E** and **1F**), this suggests a cooperative role in gene expression regulation. Partial depletion of CrsL altered the expression of numerous genes, although this effect may be underestimated due to residual CrsL protein remaining in the cells. Notably, genes affected by CrsL depletion overlap in part with those influenced by CarD depletion. Collectively, the data indicate that CrsL is a component of the mycobacterial transcriptional machinery.

In RNA-seq data, the *otsA* (*MSMEG_5892)* expression also decreased following CrsL depletion (**Figure 6A and 6B**), potentially due to dCas9 recruitment to *crsL* gene locus within the *crsL*– *otsA* operon (48,49). *otsA* encodes an enzyme in the conserved trehalose biosynthetic pathway, OtsA/B (124), and has not been reported to regulate transcription, suggesting that the differentially expressed genes identified by RNA-seq are primarily regulated by CrsL. Trehalose protects biological molecules against abiotic stresses, and OtsA has been associated with enhanced viability upon desiccation in *Rhizobium etli* (125), or the survival of *Salmonella enterica* at 50° C (126). While most prokaryotes possess only the OtsA/B pathway, mycobacteria have OtsA/B, TreY/TreZ, and TreS enzymes for trehalose synthesis (104). In *M. smegmatis*, these three pathways are functionally redundant, with trehalose levels in the Δ*otsA* mutant comparable to the wild type (103). Furthermore, Δ*otsA* shows no growth impairment at 43° C. These findings strongly indicate that the reduced growth observed upon CrsL depletion at elevated temperatures is primarily due to CrsL, not OtsA. Nevertheless, the organization of the *crsL–otsA* operon is conserved among actinobacteria, suggesting it has been maintained for its functional or regulatory advantages.

We focused specifically on the direct targets of CrsL and identified two genes linked to a temperature-sensitive phenotype: DesA (*MSMEG_5773*) is a desaturase enzyme involved in fatty acid biosynthesis, which contributes to membrane fluidity in cold stress (108,127,128). The DEAD/DEAH box helicase (*MSMEG_1930*) is an RNA chaperone that remodels RNA structures and RNA–protein complexes in an ATP-dependent manner, supporting essential processes in RNA metabolism, including transcription, RNA processing, translation and RNA decay (115,129–131). Double-stranded RNA secondary structures secondary are formed under cold stress in bacteria (132,133) and the expression of *MSMEG_1930* is highly upregulated in early cold stress in *M. smegmatis* (134). CrsL is a repressor of *DesA* and *MSMEG_1930*. CrsL binds to their promoters, and when CrsL is depleted, their expression increases. We propose that the lack of CrsL-mediated repression leads to increased expression of *DesA* and *MSMEG_1930* in the CrsL-depleted strain, which subsequently impairs the growth of *M. smegmatis* at elevated temperatures.

Besides CarD and RNAP, CrsL interacts also with the DnaK protein. DnaK (Hsp70), is an essential chaperone in *M. smegmatis* (135). Mycobacterial DnaK is involved in the native folding of important proteins, such as RNAP subunits, and also interacts with mutated, rifampicin-resistant RNAP β subunits, which increases antibiotic resistance (136). Future studies will determine whether CrsL could affect the DnaK-RNAP interaction.

As part of the Hsp70 family of heat shock proteins, *dnaK* expression increases at elevated temperatures (137). DnaK prevents protein aggregation, assists in the refolding of denatured proteins, and maintains protein quality control (138–141). In *E. coli*, DnaK regulates the availability of σ^32^, which controls the expression of heat shock response genes. Under normal conditions, DnaK associates with σ^32^, preventing σ^32^ binding to the promoters. During heat shock, accumulated denatured proteins sequester DnaK, releasing σ^32^, which then activates the expression of heat shock response genes, including *dnaK* itself. As DnaK levels increase and the amount of denatured proteins decreases, DnaK again sequesters σ^32^, shutting down σ^32^-dependent transcription (142). However, the molecular mechanisms regulating heat shock genes considerably differ among bacterial species. In mycobacteria, DnaK is synthesized from the *dnaKJE-hspR* operon, which is autoregulated by HspR repressor (137). HspR has a C-terminal hydrophobic tail, which is the primary site where DnaK binds (143). DnaK enhances the DNA-binding repressor activity of HspR (144).

CrsL has a stretch of “MLGIGP” amino acids in its sequence, which is highly conserved and can be considered as hydrophobic. In addition, CrsL appears to be an intrinsically disordered protein on its own. It is plausible that DnaK interacting with CrsL affects its folding or function. The structure of CrsL, or its interaction with DnaK (whose availability is regulated by temperature shifts and the levels of denatured proteins) could act as an additional sensor for temperature fluctuations.

Additional genes regulated by CrsL are also linked to temperature changes and protein folding. CrsL is a repressor for the *MSMEG_0024* (Rv0009 in *M. tuberculosis*) encoding PpiA, peptidyl-prolyl cis-trans isomerase B. These enzymes specifically catalyze the cis-trans isomerization of peptide bonds at proline residues, accelerating protein folding and enhancing folding efficiency. Interestingly, in *M. tuberculosis*, PpiA is repressed by HrcA (137), the second transcriptional repressor controlling heat shock genes (including *groEL2* and *groES* chaperones). *groES* and *groEL2* expression is also negatively regulated by F6 non-coding sRNA (145). F6 sRNA also negatively regulates many genes from *MSMEG_0153-MSMEG_0161*, *MSMEG_0167-MSMEG_0168*, *MSMEG_0170-MSMEG_0171* or *MSMEG_0150* (146), which were among the genes with most significantly increased expression in stationary phase after CrsL depletion (**Supplementary Table 5**). F6 sRNA is expressed from the intergenic region *MSMEG_0373-MSMEG_0374*, downstream of the *MSMEG_0373* gene (146). The expression of *MSMEG_0373* (encoding 3-ketoacyl-CoA thiolase) is directly activated by CrsL (∼1.5-fold and ∼3-fold decrease in exponential phase and stationary phase, respectively, upon CrsL depletion). This suggests that CrsL may activate the expression of F6 RNA, which subsequently negatively regulates the expression of F6 sRNA-dependent genes. Although we did not observe changes in *groEL2* and *groES* expression after CrsL depletion at optimal temperature (37° C), CrsL depletion may lead to increased expression of F6 sRNA, which could negatively regulate *groES* and *groEL2* expression at elevated temperatures. This might contribute to impaired growth of the CrsL depletion strain at 45° C.

Interestingly, the *MSMEG_0373* promoter is exclusively associated with CrsL and σ^A^, but not with RNAP, CarD and RbpA (**Supplementary Figure 4A**). The homolog of *MSMEG_0373* in *M. tuberculosis* is fadA2 (*Rv0243*). This membrane-anchored enzyme catalyzes the final step of β-oxidation in the fatty acid degradation pathway, which involves several steps to break down fatty acid molecules into acetyl-CoA, which is then utilized in the tricarboxylic acid cycle (147). FadA2 is one of six putative thiolases involved in the final step of β-oxidation. The primary σ factor, such as σ^A^, typically cannot stably bind to promoters on its own. Although we observed only CrsL and σ^A^ at the *MSMEG_0373* promoter by ChIP-seq, we cannot exclude the presence of additional proteins that might associate with CrsL and σ^A^ in a larger complex. Mycobacterial σ^A^ interacts with several transcription factors, including PhoP or CRP (cAMP receptor protein, Crp1, *MSMEG_6189*) (50), which is a transcriptional regulator that controls gene expression by recognizing altered cAMP levels in bacteria. We propose that there may be alternative, noncanonical mechanisms for regulating σ^A^-dependent transcription in mycobacteria, which are rare, but still present and should be considered.

Similarly to CrsL, many proteins have been identified as being intrinsically disordered and unable to form tertiary structures under normal conditions over the past three decades (148,149). Despite lacking a defined secondary structure, these proteins remain functional and interact with various partners, serving as hub proteins that facilitate molecular communication through protein-protein interactions (150,151). Moreover, many disordered proteins exist in a transient state and have some sort of preformed structural motifs (152,153). These proteins can also adopt a defined fold upon binding with other protein(s) both in prokaryotes (152,154) and eukaryotes (155–157). In CrsL, two regions (aa 20-27 and 34-48) are predicted to be potentially structured. A CrsL-CarD interaction predicted by AlphaFold Multimer showed that these regions might be folded (**Supplementary Figure 3B, 3C and 3E**). Future experiments will be required to address their importance. The unstructured nature of CrsL then might enable its rapid association with target protein(s) (swift binding) and allow it to flexibly interact with diverse binding partners, and might also make CrsL well-suited for sensing and responding to temperature-based shifts.

In conclusion, this work identifies CrsL as a mycobacterial transcription factor and provides a foundation for future studies of this protein focusing on molecular details of its interaction with CarD/RNAP, its mechanistic role, and its impact on cell physiology, including the regulation of genes associated with temperature changes.

## DATA AVAILABILITY

Sequencing data are available at ArrayExpress: *M. smegmatis* RNA-seq for *carD* and *crsL* knockdown [E-MTAB-13812], *M. smegmatis* CrsL ChIP-seq [E-MTAB-12348]. All original code has been deposited on Zenodo under DOI:10.5281/zenodo.11174175. Our webpage (msmegseq.elixir-czech.cz) can be used for visualizing all ChIP-seq and RNA-seq datasets. The mass spectrometry proteomics data have been deposited to the ProteomeXchange Consortium via the PRIDE partner repository with the dataset identifier PXD058166.

## AUTHOR CONTRIBUTIONS

Mahmoud Shoman: Investigation, Methodology, Visualisation, Writing – Original Draft; Jitka Jirát Matějčková: Investigation, Methodology; Marek Schwarz: Methodology, Bioinformatics Analysis, Data Curation, Software; Martin Černý: Investigation, Methodology, Nabajyoti Borah: Investigation, Viola Vaňková Hausnerová: Conceptualization, Methodology; Michaela Šiková: Investigation; Petr Halada: Investigation; Hana Šanderová: Conceptualization, Methodology; Petr Halada: Investigation; Martin Hubálek: Investigation; Věra Dvořáková: Investigation; Martin Převorovský: Methodology, Data Curation; Jana Holubová: Investigation, Methodology; Ondřej Staněk: Investigation, Methodology Libor Krásný: Funding Acquisition, Data Discussion;; Lukáš Žídek: Funding Acquisition, Supervision; Jarmila Hnilicová: Funding Acquisition, Supervision, Visualisation and Data Discussion.

## CONFLICT OF INTEREST STATEMENT

None declared.

## ACKNOWLEDGMENTS

We acknowledge Josef Dadok National NMR Centre of CIISB, Instruct-CZ Centre, supported by MEYS CR (LM2023042) and European Regional Development Fund-Project „Innovation of Czech Infrastructure for Integrative Structural Biology“ (No. CZ.02.01.01/00/23_015/0008175) for the NMR measurement time. The RNA spike was kindly gifted from Dr. Radek Malík, IMG, Prague.

## FUNDING

Czech Science Foundation [23-05622S J. Hn. and No. 22–12023S to L. Ž.]; The Charles University Grant Agency [275823 to M. Sh.]; European Union – Next Generation EU, National Institute of Virology and Bacteriology [EXCELES LX22NPO5103 to N.B.]; The Ministry of Education, Youth and Sports of the Czech Republic [OPJAK project CZ.02.01.01/00/22_008/0004597 to O.S. and P.H., CZ.02.01.01/00/22_008/0004575 to L.K and H.Š., research infrastructure projects ELIXIR CZ, No. LM2023055 to M. Sch]; The Ministry of Health [The National Institute of Public Health (NIPH), 75010330 to V.D.].

## Notes

### Competing Interest Statement

The authors have declared no competing interest.

